# A CellAgeClock for expedited discovery of anti-ageing compounds

**DOI:** 10.1101/803676

**Authors:** Celia Lujan, Eleanor J. Tyler, Simone Ecker, Amy P. Webster, Eleanor R. Stead, Victoria E. Martinez Miguel, Deborah Milligan, James C. Garbe, Martha R. Stampfer, Stephan Beck, Robert Lowe, Cleo L. Bishop, Ivana Bjedov

## Abstract

We aim to improve anti-ageing drug discovery, currently achieved through laborious and lengthy longevity analysis. Recent studies demonstrated that the most accurate molecular method to measure human age is based on CpG methylation profiles, as exemplified by several epigenetics clocks that can accurately predict an individual’s age. Here, we developed CellAgeClock, a new epigenetic clock that measures subtle ageing changes in primary human cells *in vitro*. As such, it provides a unique tool to measure effects of relatively short pharmacological treatments on ageing. We validated the CellAgeClock against known longevity drugs such as rapamycin and trametinib. Moreover, we uncovered novel anti-ageing drugs, torin2 and Dactolisib (BEZ-235), demonstrating the value of our approach as a screening and discovery platform for anti-ageing strategies. The CellAgeClock outperforms other epigenetic clocks in measuring subtle ageing changes in primary human cells in culture. The tested drug treatments reduced senescence and other ageing markers, further consolidating our approach as a screening platform. Finally, we show that the novel anti-ageing drugs we uncovered *in vitro*, indeed increased longevity *in vivo*. Our method expands the scope of CpG methylation profiling from measuring human chronological and biological age from human samples in years, to accurately and rapidly detecting anti-ageing potential of drugs using human cells *in vitro*, providing a novel accelerated discovery platform to test sought after geroprotectors.

---

One of the remarkable achievements of developed countries is a continuous increase in life expectancy at birth, leading to greater longevity. However, a higher proportion of elderly in modern societies is accompanied by a steep increase in people suffering from age-related diseases. For example, cancer incidence rates, currently at 17 million worldwide, are expected to increase to 26 million in 2040 (Wilson et al. 2019), and a similar rise is expected for Alzheimer’s and Parkinson’s disease (Reeve et al. 2014). Compression of late-life morbidity is, therefore, a priority to alleviate suffering in the elderly (Partridge et al. 2018) and to reduce a growing economic burden to society (Rae et al. 2010).

Critically, seminal discoveries in the biology of ageing showed that ageing is a malleable process and that down-regulation of major cellular nutrient signalling pathways, either glucose-sensing insulin signalling or amino acid-sensing target-of-rapamycin signalling, results in longevity and health improvement in all model organisms tested from yeast to mammals (Lopez-Otin et al. 2013). For instance, the long-lived mutants in *C. elegans* are protected from tumorous cell proliferation (Pinkston et al. 2006) and have reduced toxic protein aggregation (Cohen et al. 2006), while *Drosophila* show less deterioration in their hearts (Wessells et al. 2004). Long-lived mouse mutants are protected from osteoporosis, cataracts and skin pathology, as well as decline in glucose homeostasis, immune and motor function (Selman et al. 2008). The effect of these mutations is conserved from yeast to mammals, and it is, therefore, expected that if drugs replicate the biological impact of these changes, this could improve health in the elderly and prevent age-related disease. It is increasingly recognised that directly targeting ageing through pharmacological interventions, as opposed to specific age-related diseases, is a highly promising strategy for broad-spectrum disease protection (Niccoli and Partridge 2012). However, at present, there are only a handful of reliable anti-ageing drugs whose effects have been confirmed in mammals, such as rapamycin (Harrison et al. 2009) and metformin (Martin-Montalvo et al. 2013). Crucially, there are currently no sufficiently reliable ageing biomarkers for testing drugs on human cells *in vitro*, and the development of a specialised epigenetic clock is a promising approach (Castillo-Quan et al. 2015; Field et al. 2018; Horvath et al. 2018; Bell et al. 2019; Horvath et al. 2019).

To accelerate the discovery workflow for anti-ageing drugs, we took advantage of the breakthrough in the ageing field which showed that epigenetic clocks provide the most accurate measurements of human age. For instance, the approximate error rate for the Skin and Blood clock is ±2.5 years (maximal correlation coefficient 0,98) (Horvath et al. 2018). Epigenetic clocks surpass the accuracy of other ageing biomarkers such as telomere length and those based on transcriptomic, metabolomic or proteomic approaches, potentially because the latter approaches detect more transient and less stable cellular changes (Horvath 2013). Ageing is accompanied by overall CpG hypomethylation, whilst some CpG islands and gene regions become hypermethylated (Booth and Brunet 2016). Remarkably, only a small selection of the 56 million CpG sites in the diploid human genome, coupled with computational algorithms, is sufficient to provide an accurate readout of human age. One of the first epigenetic clocks was developed by Hannum using just 71 CpG sites to estimate age from blood samples (Hannum et al. 2013), while Horvath’s multi-tissue age estimator (Horvath 2013) and Skin and Blood clock (Horvath et al. 2018) use 353 and 391 CpG sites, respectively (Field et al. 2018; Horvath and Raj 2018). Even a single CpG site in the ELOVL2 gene is sufficient to determine age (Garagnani et al. 2012), albeit clocks using only a few CpG sites are less accurate and less applicable to different tissues (Horvath and Raj 2018). The epigenetic clocks measure the ageing process inherent to all our cells and tissues, irrespective of their proliferation rate (Horvath et al. 2019). As the human epigenome reflects physiological changes, epigenetic clocks cannot only predict chronological age from a human sample but also give an estimate of biological age as has widely been demonstrated by the associations of epigenetic age with morbidity and mortality (Marioni et al. 2015; Horvath and Raj 2018). Recently, valuable predictors focussing on this aspect have been developed: PhenoAge (Levine et al. 2018) and GrimAge (Lu et al. 2019), which form the best epigenetic morbidity and mortality predictors available to date.

DNA methylation also captures information on the approximate number of cell divisions a cell has been through, as has been shown by epiTOC (Yang et al. 2016), a mitotic-like clock that approximates stem cell divisions and correlates with cancer risk (Tomasetti et al. 2017), and MiAge, which also measures mitotic age (Youn and Wang 2018). The biology underlying CpG methylation alterations at the sites linked to ageing clocks is not well understood. The exception is ribosomal clock based on CpG methylation in ribosomal RNA (rRNA), which is highly conserved throughout evolution and which forms nucleolus that has itself been implicated in ageing (Tiku and Antebi 2018; Wang and Lemos 2019). Horvath suggests an interesting hypothesis that epigenetic maintenance programmes are being reflected in DNA methylation alterations (Horvath 2013; Horvath and Raj 2018; Raj and Horvath 2020). Recent findings implicate loss of H3K36 histone methyltransferase NSD1 in epigenetic ageing clock acceleration (Martin-Herranz et al. 2019). Despite the enigma regarding the mechanism of epigenetic clocks, they are reliable predictors of age and extremely useful biomarkers (Field et al. 2018; Horvath and Raj 2018; Bell et al. 2019). However, little is known so far about the performance of these clocks in *in vitro* ageing experiments. It has recently been shown that the rate of epigenetic ageing in cultured cells is significantly faster than in the human body (Horvath et al. 2019; Sturm et al. 2019) and that epigenetic age is retarded by rapamycin *in vitro* (Horvath et al. 2019), but neither of the clocks specialised for *in vitro* drug discovery nor were they tested on multiple anti-ageing drugs.

Therefore, we aimed to exploit the exceptional accuracy of CpG methylation clocks to uncover new anti-ageing pharmacological treatments. The current gold standard for discovering novel anti-ageing drugs are longevity experiments, which are laborious, lengthy and expensive. For instance, in mice, they take three years, thereby precluding any large scale drug screens. Existing screens in *C. elegans* commonly use live *E. coli* as food (Lucanic et al. 2013; Ye et al. 2014), which is a disadvantage as drugs are metabolised first by the bacteria making their effect on worms secondary, which may lead to confounded results (Cabreiro et al. 2013; Pryor et al. 2019). Yeast drug screens lack the crucial aspect of tissue toxicity (Zimmermann et al. 2018). In addition, all longevity assays require constant supply of the drug, making them highly expensive. Other attempts to uncover anti-ageing effects of drugs are based on computational analysis using existing transcriptomic information on the ageing process combined with drug characteristics (Donertas et al. 2018). However, transcriptomic changes are more transient and noisy when compared to DNA methylation and are, therefore, a less consistent ageing marker (Horvath and Raj 2018).

We tested if existing epigenetic clocks could be used to measure anti-ageing drug potential in human primary cells *in vitro* and if we could build a new clock specialised for this purpose. Senescence is tightly associated with ageing of the organism, and because of the pronounced resemblance of ageing in primary cells *in vitro* to ageing *in vivo*, together with the evidence that human DNA methylation signatures are conserved and accelerated in cultured fibroblasts (Sturm et al. 2019), we used cultured human cells as a proxy for human ageing (Lowe et al. 2015; Horvath et al. 2019). The ability to test anti-ageing drug properties directly on human cells *in vitro* could considerably accelerate the discovery of new compounds promoting healthy ageing. To this end, we used normal human mammary fibroblasts (HMFs) from a healthy 16-year old donor that we cultured from passage 10 to passage 20, which is before these cells reach senescence at passage 29 (Supplemental Fig. S1A-D). To measure CpG methylation, we used EPIC Arrays (Illumina) that measure methylation at 850,000 sites.

First, we tested the three most suitable existing epigenetic clocks, to determine if they could detect weekly and monthly ageing differences occurring during serial passaging of HMFs (Fig. 1A). The Multi-tissue clock (Horvath 2013) consistently predicted a higher epigenetic age, and at passage ten this was 43.6±1.0 years (Fig. 1A), consistent with what was recently reported (Sturm et al. 2019). This increased age estimate, compared to the age of the donor who was 16 years old, is in accordance with published data demonstrating that this epigenetic clock overestimates the age of mammary tissue samples (Horvath 2013). The PhenoAge clock (Levine et al. 2018), developed to predict mortality and morbidity risks, reported the epigenetic age of the donor to be 3.5±1.1 years (Fig. 1A). The most accurate age estimate, predicting the age of the donor at 23.2±0.87 years, was obtained using the Skin and Blood clock, which is specialised for determining donor age of easily accessible human tissues and cells in culture (Fig. 1A). The Multi-tissue clock and Skin and Blood clock showed a small increase in age with progressive passaging (from passage 10 to 20, age estimate increased from 43.6±1.0 to 53.9±1.7 and from 23.2±0.87 to 31.6±1.2 years, respectively), while this increase was greater for the PhenoAge clock (from 3.5±1.1 to 26.6±9.7 years). This suggested that, of the tested clocks, the PhenoAge clock captures ageing *in vitro* best (Fig. 1A). However, the PhenoAge clock showed substantial variability in predictions for higher passages, which would obstruct the detection of subtle ageing differences upon anti-ageing drug treatments. In conclusion, while the Skin and Blood clock (Horvath et al. 2018) measures fibroblast ageing in culture, none of the existing clocks was ideally suited to accurately measure subtle anti-ageing drug potential in human primary cells *in vitro*, and similar comparisons have recently been reported by others (Horvath et al. 2019; Sturm et al. 2019).

**Figure 1.**
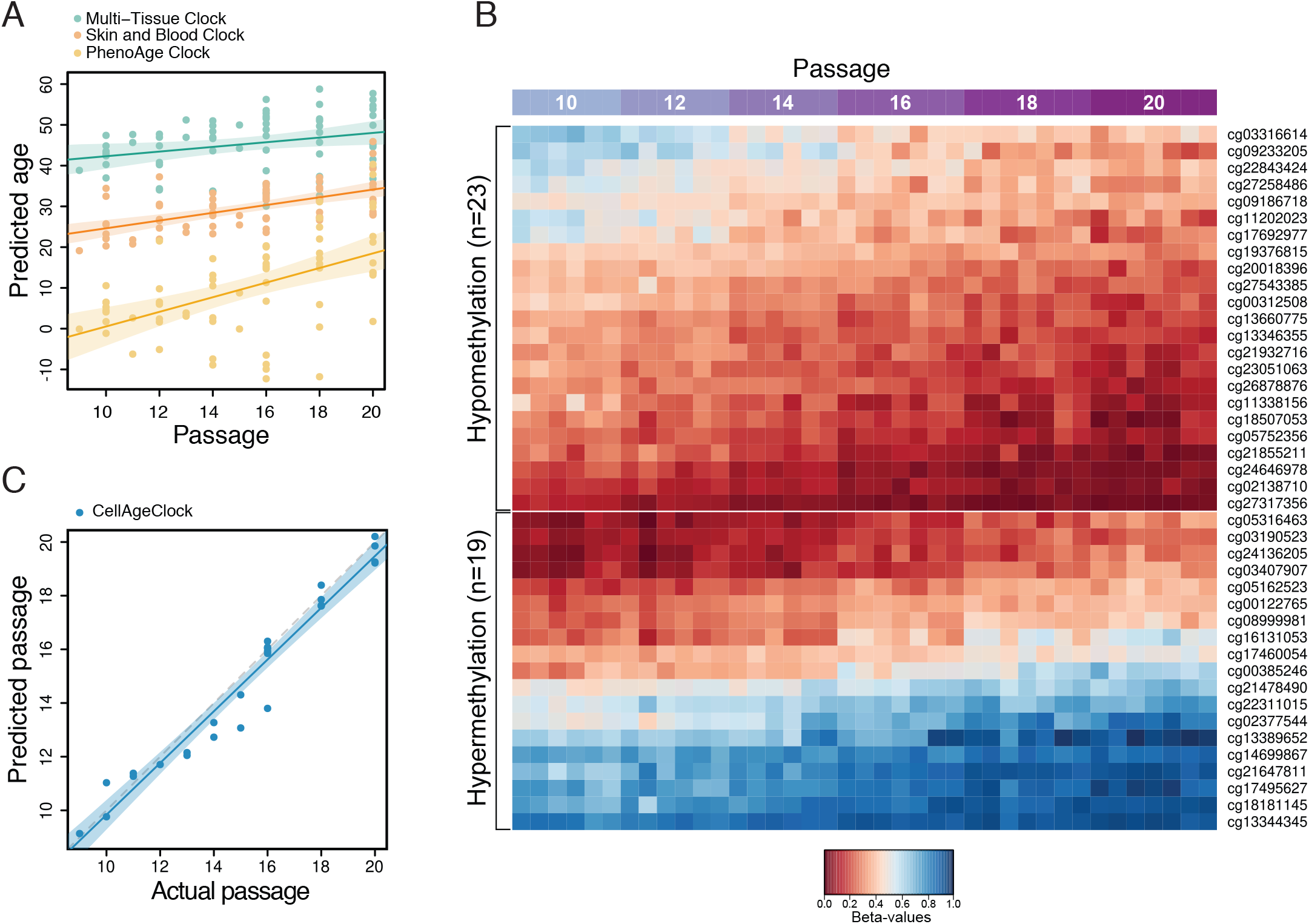
Development of CellAgeClock for monitoring subtle ageing difference in cells in culture. A) Predicted age of control samples using three existing epigenetic clocks. Predicted epigenetic age for control samples across all experiments as estimated by the Multi-tissue clock (green), the Skin and Blood clock (orange) and the PhenoAge clock (yellow). Fitted lines are shown with 95% confidence intervals (semi-transparent). All three clocks show a trend to increase in predicted age with progressing passage, however there is considerable variability in predictions, particularly for the PhenoAge clock. The Multi-tissue clock consistently predicted cells to have the highest epigenetic age, while the PhenoAge clock consistently predicted cells to have the lowest epigenetic age, which even reached below zero for several samples at various passages. B) Heatmap representing 23 CpG probes that undergo hypomethylation with increasing cell passage and 19 CpGs that undergo hypermethylation with increasing cell passage. These 42 CpG probes were used to develop the CellAgeClock. Probes that undergo hypomethylation and hypermethylation with increasing passage were separated and ordered by their methylation values per row. The mean absolute difference between passage 20 and passage 10 among clock CpGs is 0.2. C) Testing the CellAgeClock on HMF and HDF samples that were not used to train the clock. The gray dashed line represents the diagonal (perfect prediction). The fitted line of the actual data is shown in blue, with a 95% confidence interval (semi-transparent). Cell passages are predicted accurately.

This prompted us to develop a new clock that, rather than predicting donor age in years, specialises in measuring methylation changes occurring during ageing of primary cells in culture and could differentiate DNA methylation state between each passage. To this end, we developed a clock using two different cell types, the above-mentioned HMFs and human dermal fibroblasts (HDFs), which were obtained from a different donor, have a different proliferative lifespan *in vitro*, and a different rate of DNA methylation change. Like the HMFs, the HDFs were serially passaged and sampled every other passage for DNA methylation analysis.

We used a total of 39 HMF and HDF samples to build the clock (see Materials and methods). To preselect informative probes, we performed a statistical test to identify CpGs undergoing DNA methylation changes with increasing cell passage using linear regression (Supplemental Fig. S2A). The resulting 2,543 CpGs were used to build the clock model by elastic net regression, similar to the method used by Horvath (Horvath 2013). The model selected 42 predictor CpGs (“clock CpGs”), shown in Fig. 1B and Supplemental Fig. S2B. Of these CpGs, 23 undergo hypomethylation and 19 become hypermethylated with increasing cell passage (Fig. 1B and Supplemental Fig. S2B). Sixteen of the CpGs are located in intergenic regions (IGRs), whereas 14 of them are located in gene bodies and 12 in promoters, respectively (Supplemental Fig. S2C). Interestingly, two of the clock CpGs map to gene *GRID1*, one is located in its 3’UTR and one in the gene’s body. *GRID1* encodes a subunit of glutamate receptor channels. Several other clock CpGs are also located in genes implicated in cell receptor activity and metabolic processes, such as *LDLRAD4 and NPSR1*. Furthermore, multiple clock CpGs map to genes that play roles development as well as in the regulation of transcription and protein binding. Examples include *GGN, MEIS2, NF1, PROP1, RFX4, RUNX3* and *SMARCA2*. The 42 CpGs together with their detailed genomic and functional annotation are available in Table 1.

After building our novel epigenetic clock to measure cell ageing *in vitro*, named CellAgeClock, we tested its performance using an entirely different set of samples (n=26), consisting of 22 HMF and four HDF samples. We observed accurate prediction of passage number for both HMFs and HDFs, with a Root Mean Square Error (RMSE) of 0.37 (Fig. 1C). To compare the performance of the CellAgeClock with other epigenetic age predictors, we calculated Spearman’s rank correlation coefficients between the clocks’ output and actual cell passage (see Table 2). The CellAgeClock showed the best correlation among the tested predictors, with Spearman’s Rho = 0.98 and p < 2.2e-16. We also tested the mitotic-like clocks EpiTOC and MiAge, for comparison. However, their correlation coefficients were negative, small and non-significant (Rho > −0.3, p > 0.05).

Having built a precise epigenetic clock that measures methylation changes during replicative ageing of human primary cells *in vitro*, we tested if anti-ageing drug treatment of HMFs and HDFs decelerated the CellAgeClock. We chose an mTOR inhibitor, rapamycin, which is one of the most robust and evolutionarily conserved anti-ageing drug targets (Saxton and Sabatini 2017), and which mediates its effect through down-regulation of S6K and Pol III, and up-regulation of autophagy (Bjedov et al. 2010; Filer et al. 2017). We chose relatively low rapamycin concentration of 5nM that did not inhibit cell growth (Supplemental Fig. S1A) but moderately downregulated mTOR signalling, as evidenced by decreased pS6K and p4E-BP phosphorylation (Supplemental Fig. S3). This setup mimics the pro-longevity effects of rapamycin *in vivo* where it is well accepted that only mild nutrient sensing pathway inhibition increases life- and healthspan (Bjedov and Partridge 2011; Lopez-Otin et al. 2013).

DNA methylation profiles from HDF and HMFs collected following four, six and eight weeks of rapamycin treatment (passage 16, 18 and 20; Fig. 2) were analysed using the CellAgeClock and clearly demonstrated that rapamycin slows down methylation changes associated with replicative ageing. Interestingly, this clock deceleration was more pronounced upon longer treatment as shown by the gradual decrease of predicted-actual passage from 16 to 20 weeks. The low dose rapamycin treatment did not affect population doublings, confirming that the methylation changes were not a reflection of proliferation inhibition or slowing of the cell cycle (Supplemental Fig. S1). This is further evidenced by comparing the predicted passage from the CellAgeClock against cumulative population doubling, showing that rapamycin samples lie on a separate line to that of the control samples (Supplemental Fig. S4A-B). Contrarily, rapamycin samples and controls differed to considerably lesser extent when actual passage and cumulative population doublings are compared (Supplemental Fig. S4A-B). We observed a similar pattern for HMFs and HDFs (Fig. 2), suggesting that the CellAgeClock is applicable to different cells, albeit calibration is required for cells that reach senescence at different rates.

**Figure 2.**
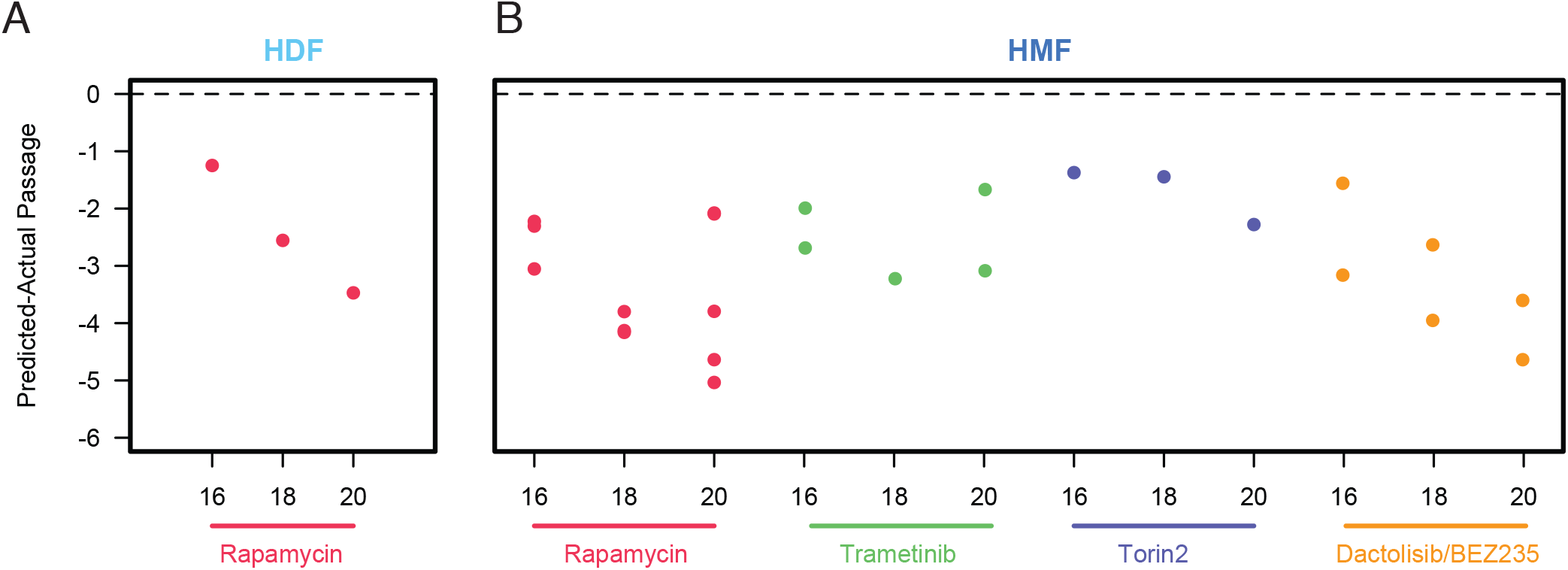
Using the CellAgeClock for the detection of anti-ageing drugs. A) The CellAgeClock predictions of Human Dermal Fibroblasts (HDF) and B) Human Mammary Fibroblasts (HMF). Represented is Predicted-Actual Passage for Passage 16, 18 and 20, showing deceleration of the CellAgeClock upon treatment with anti-ageing drugs rapamycin (5nM), Dactolisib/BEZ235 (10nM), torin2 (5nM) and Trametinib (0.1nM).

We then focused on HMFs to test another anti-ageing drug, trametinib (Slack et al. 2015), an inhibitor of the MEK/ERK signalling pathway, which we also applied in low concentration to avoid any effect on growth and population doubling (Supplemental Fig. S1B and S3). The CellAgeClock analysis of trametinib treatment showed clock deceleration for all three passages tested (Fig. 2), thereby confirming previous results in *Drosophila in vivo* that trametinib extends lifespan (Slack et al. 2015). Next, we examined the effect of two other inhibitors of nutrient-sensing pathways as mutations in these pathways in model organisms represent the most evolutionary conserved anti-ageing interventions (Lopez-Otin et al. 2013). We tested Dactolisib/BEZ235, a dual ATP competitive PI3K and mTOR inhibitor, for which we again optimised the dose of the treatment to obtain a reduction in signalling, as shown by pS6K downstream target 4E-BP (Supplemental Fig. S3), without significant proliferation impairment (Supplemental Fig. S1C). Dactolisib/BEZ235 slowed down the DNA methylation changes similar to rapamycin, suggesting that Dactolisib/BEZ235 could be a new anti-ageing drug according to the output of the CellAgeClock (Fig. 2). We also tested torin2, which is a selective inhibitor of the mTOR pathway that inhibits both mTORC1 and mTORC2, unlike rapamycin, which targets solely mTORC1. Owing to its more complete inhibition of the mTOR pathway, we were interested in examining its effect on replicative ageing, especially as the role of mTORC2 in ageing is less well established. The impact of mTORC2 inhibition on lifespan can be positive or negative depending on which of the mTORC2 downstream effectors is affected, in which tissue, and whether females or male mice are used for the experiment (Kennedy and Lamming 2016). Some of the negative effects of mTOR pathway inhibition, such as insulin resistance and hyperlipidemia, are attributed to the mTORC2 branch of the pathway and may arise under certain conditions of prolonged and/or high dose rapamycin treatment (Kennedy and Lamming 2016). Interestingly, while our CellAgeClock suggests that torin2 is indeed a novel anti-ageing drug (Fig. 2, Supplemental Fig. S1D and S3), its effect on ageing in mammalian cell culture appears to be less pronounced than that of rapamycin. This is in line with literature suggesting that a promising strategy to improve healthy ageing is the development of inhibitors that are highly specific for mTORC1 or that target mTORC1 downstream effectors separately (Kennedy and Lamming 2016).

Next, we compared our anti-ageing drug screening results obtained by the CellAgeClock with analyses using Horvath’s Multi-tissue and Skin and Blood clock, as well as the PhenoAge clock. The clocks did not detect any significant effect of anti-ageing drug treatment (Supplemental Fig. S5). The Skin and Blood clock^26^ was used recently to measure deceleration of ageing in primary fibroblasts (Horvath et al. 2019; Sturm et al. 2019), however the concentration of rapamycin used in our conditions was five times lower without effect on cell growth, highlighting the sensitivity of our epigenetic clock to detect age-related methylation changes at very low drug concentrations. Under our conditions, the only epigenetic clock that detected gradual methylation changes from passage 10 to passage 20 was the PhenoAge clock (Supplemental Fig. S5). However, its output was more variable between samples and inconsistent for anti-ageing drug treatments, reporting both clock acceleration and deceleration. For instance, rapamycin, Dactolisib/BEZ235 and torin2 treated cells appeared slightly younger compared to controls, whereas trametinib treated cells were estimated older to some extent (Supplemental Fig. S5), unlike the results we obtained with our CellAgeClock (Fig. 2). Overall, the CellAgeClock that we developed here was more consistent and performed significantly better on ageing cells in culture and following known anti-ageing drug treatments compared to existing clocks. Our results are supportive of clocks being highly specialised for a certain task, and suggests that while other popular epigenetic clocks perform remarkably on determining donor’s age in years and their health status, they were not able to robustly detect slight ageing changes in human primary cells induced by drug treatment over a short period of time *in vitro*.

Next, we assessed if the CellAgeClock is suitable for the screening of novel anti-ageing drugs. To this aim, we examined if drugs that decelerate the CellAgeClock also reduce features associated with senescence, such as morphological changes and expression of ageing biomarkers (Hanzelmann et al. 2015). Rapamycin, Dactolisib/BEZ235 and Trametinib treatment slowed down morphological alteration in cells that gradually occur during replicative ageing, namely cell elongation, increased nuclear area and cell area, and the treated cells appeared particularly ‘youthful’ (Fig. 3). Another characteristic of senescence is increased expression of the cyclin-dependent kinase inhibitors p21^CIP1/Waf1^ and p16^INK4a^. p21^CIP1/Waf1^ triggers G1 cycle arrest upon damage and can lead to senescence or apoptosis (He and Sharpless 2017; McHugh and Gil 2018). Expression of p16^INK4a^, which is produced from the CDKN2A gene together with p19^ARF^ (p14^ARF^ in humans) increases exponentially during ageing and was suggested to stabilise the senescent state (Gire and Dulic 2015). p16^INK4a^ expression was the marker of choice for senescent cell clearance leading to prolonged lifespan in mice (Baker et al. 2016).

**Figure 3.**
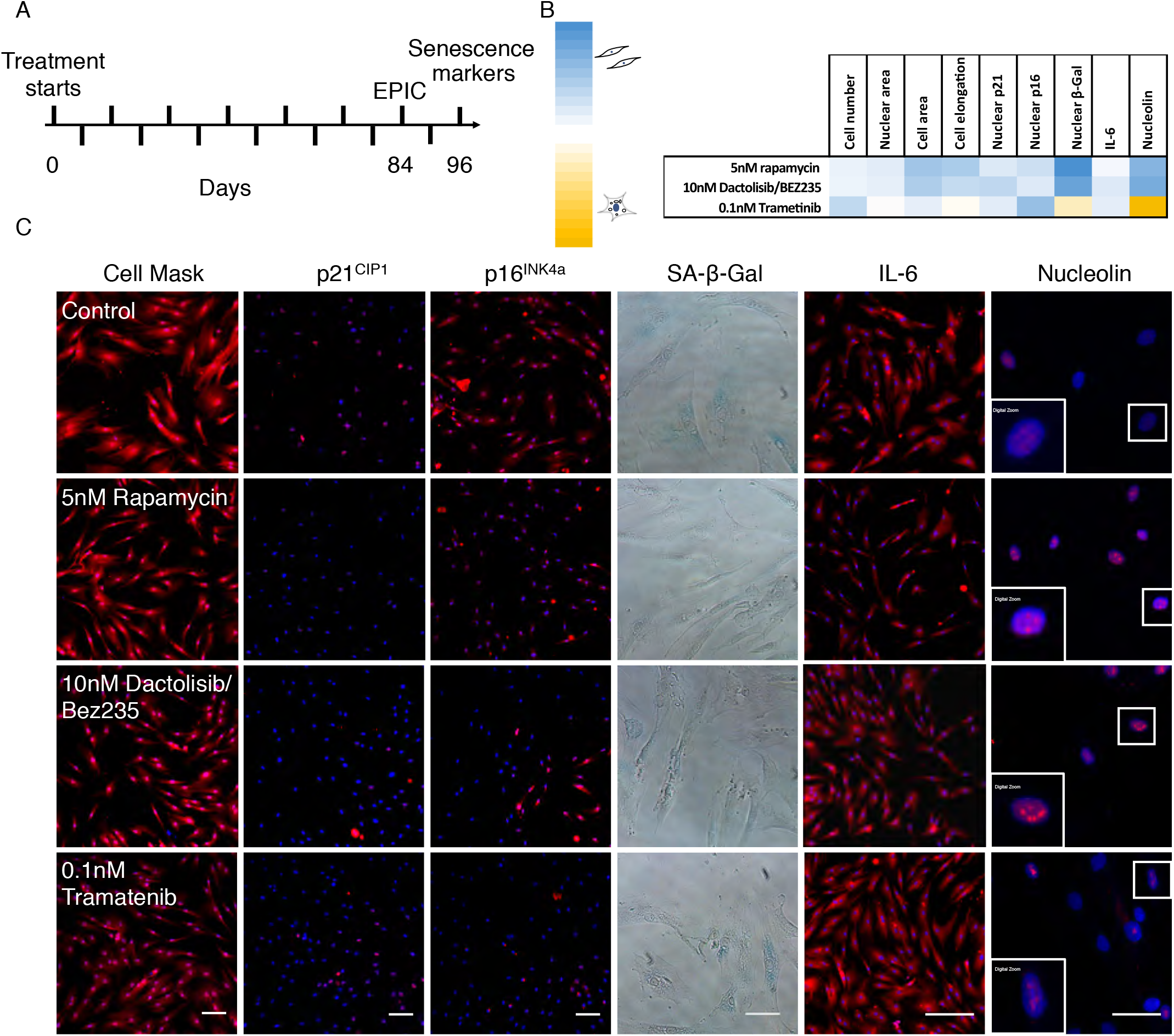
Treatment with anti-ageing drugs decreases markers of senescence. A) Schematic illustrating the experimental set-up conducted in P10 to P22 HMFs, passaged weekly. B) Multi-parameter analysis of senescence markers. Colour coding used to illustrate the number of Z scores of the experimental drug value from the respective control mean. Scores highlighted in red denote a shift towards a more proliferative phenotype and scores highlighted in green denote a shift to a more senescent phenotype. C) P22 HMFs stained with DAPI (blue) and Cell Mask, p21, p16, IL-6, or nucleolin (red), or SA-β-Gal (blue) following 96-day treatment with 5nM Rapamycin, 10nM Dactolisib/BEZ235, 0.1nM Trametinib or their respective controls. Size bar, 100*μ*m.

Our results demonstrate that drugs which decelerate the CellAgeClock at the same time reduce expression of both nuclear p21^CIP1/Waf1^ and p16^INK4a^ compared to non-treated cells, showing their efficacy in delaying the senescence programme (Fig. 3B-C). In addition, the most frequently used senescent marker, senescent-associated β-galactosidase activity (SA-βgal), was significantly decreased upon anti-ageing drugs treatment with rapamycin and Dactolisib/BEZ235, but not in cells treated with trametinib (Fig. 3B-C). Another difference in senescent markers was observed with interleukin-6 (IL-6), which is one of the most important inflammatory cytokines and part of the senescent-associated secretory phenotype. IL-6 was significantly reduced in aged cells upon rapamycin and Dactolisib/BEZ235 treatment but not in trametinib treated cells (Fig. 3B-C). This difference possibly stems from the overactivated RAS/ERK pathway being a more prominent inducer of senescence than the overactivated mTOR/PI3K pathway (Kennedy et al. 2011), and hence corresponding inhibitors have different potency in inhibiting senescence. Finally, we examined the nucleolus, an organelle dedicated to rRNA production and ribosomal assembly, as it has recently emerged that maintenance of its structure, and low levels of nucleolar methyltransferase fibrillarin, is a common denominator for major anti-ageing intervention from worms to mice (Tiku and Antebi 2018). We observed that as a consequence of ageing, nucleoli in aged HMFs lose their defined round shape, are more diffused, and stain less well. For rapamycin and Dactolisib/BEZ235, we observed clearly defined and ‘younger’ looking nucleoli in aged cells. However, trametinib treated cells resembled the nucleoli of controls. In summary, a panel of the most frequently used markers for cell senescence confirmed that drugs which decelerate the CellAgeClock also make the cells appear more youthful. This strongly suggests that the CellAgeClock can be used as a robust and sensitive detector of novel anti-ageing treatments.

Finally, having discovered two novel potential anti-ageing drug treatments using the CellAgeClock, Dactolisib/BEZ235 and torin2, we tested and validated them *in vivo* using the fruit fly *Drosophila melanogaster* as a model organism. This is important as tissue-specific drug toxicity, which can be missed in cell culture, is one of the major reasons for drug failure in clinical trials. For *in vivo* longevity studies we used the outbred wild-type *w*^*Dah*^ strain which is particularly suitable for ageing studies, Drosoflipper device for fast fly transfer, and specially formulated holidic medium (Piper et al. 2014) to increase drug bioavailability compared to standard sugar-yeast-agar fly food. We used rapamycin as a positive control for longevity experiments in flies and showed that median lifespan extension on holidic media varied from 7% to 9% compared to ethanol solvent control, depending on 1*μ*M or 5*μ*M concentration, respectively (p<0.001, log-rank test), which is comparable to published literature (Fan et al. 2015) (Fig. 4A). Importantly, both Dactolisib/BEZ235 and torin2 significantly extended lifespan in *Drosophila* for 7% (p<0.001, log-rank test) (Fig. 4B-C). This firmly demonstrates that drugs that decelerate the CellAgeClock have similarly favourable output on major anti-ageing biomarkers *in vitro* and extend longevity *in vivo* (Fig. 4D).

**Figure 4.**
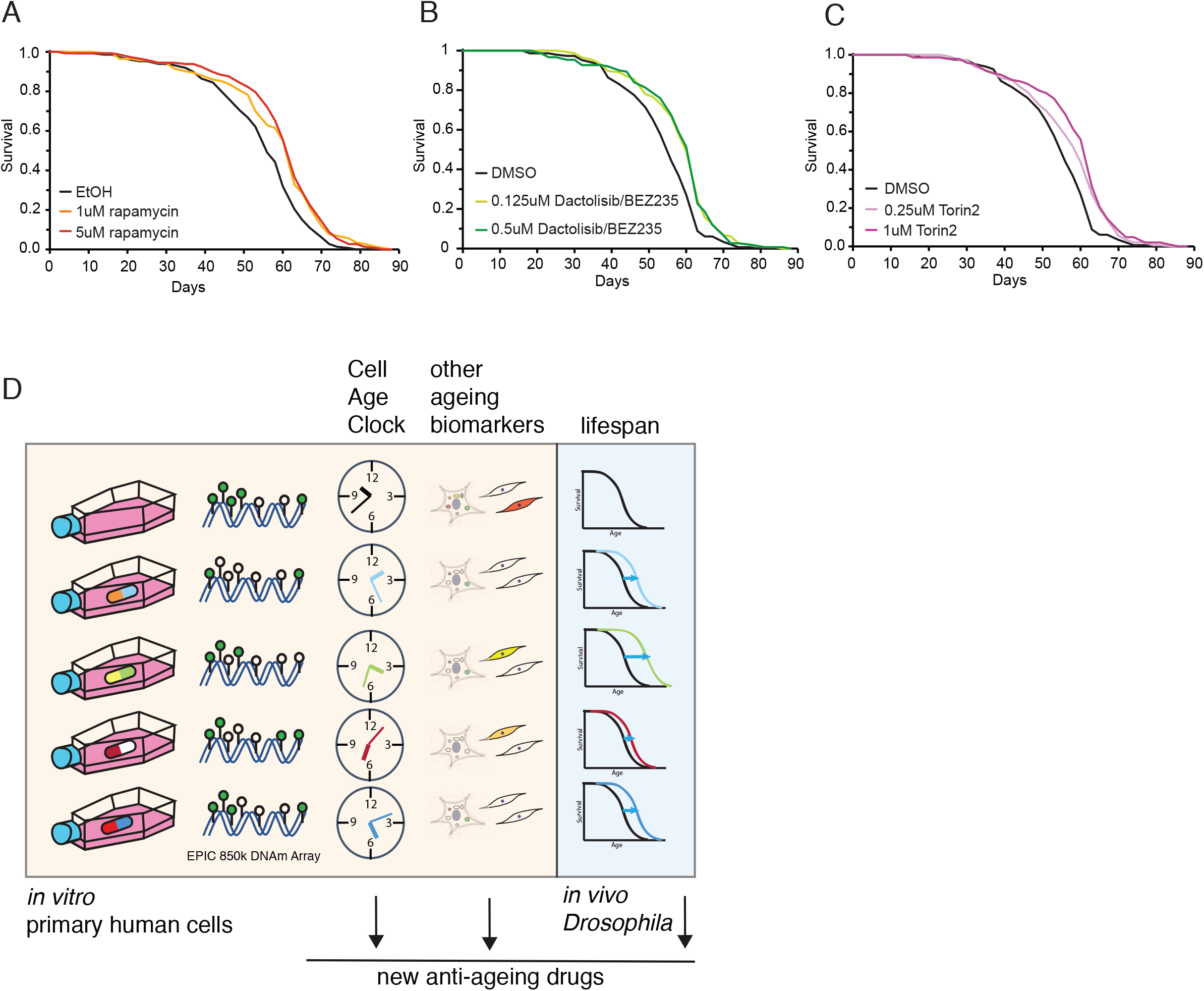
Drugs that decelerate the CellAgeClock extend lifespan *in vivo*. A) Lifespan analysis on *w*^*Dah*^ background wild type flies fed with SYA food containing different concentration of rapamycin or ethanol as solvent control. For each condition, 150 flies were used. B) Lifespan analysis on *w*^*Dah*^ background wild type flies fed with SYA food containing different concentration of Dactolisib/BEZ235 or DMSO as solvent control. For each condition, 150 flies were used. C) Lifespan analysis on *w*^*Dah*^ background wild type flies fed with SYA food containing different concentration of torin2 or DMSO as solvent control. For each condition, 150 flies were used. D) Schematic representation of our approach combining novel CellAgeClock and other ageing biomarkers *in vitro* primary human cell, together with *in vivo Drosophila* lifespan experiments, for a detailed and robust capture of anti-ageing drug potential.

For the first time, we have a robust epigenetic clock for the rapid discovery of anti-ageing drugs directly in human cells, bypassing lower model organisms and significantly shortening discovery time compared to 3-year long mice longevity analysis. Testing different compounds for ageing using the CellAgeClock could potentially reveal new anti-ageing pathways and help us to improve our knowledge base of not only ageing biology but of molecular pathways underpinning the epigenetic clocks as well, understanding of which is limited. Other available epigenetic clocks could not accurately detect the effect of low-dose short-term anti-ageing drugs on cells *in vitro*. We expect many biological outputs to be extracted by the CellAgeClock and other epigenetic clock algorithms in the future, given the wealth of information stored in our epigenome.

Better experimental systems to test anti-ageing drugs are very much needed, given a rising proportion of the elderly in modern societies and, as a consequence, larger numbers of people suffering from age-related diseases. Our results show that by using the CellAgeClock, cultured primary human cells can be used as a proxy to measure human ageing and can reliably detect anti-ageing effects upon a relatively brief treatment. By doing so, this fast and accurate method is expected to accelerate the discovery of novel preventive treatments for age-related disease, directly using human cells. Importantly, follow-up research will be focused on expanding our findings on different types of primary cells from donors of different ages, as well as on testing further compounds. While ageing itself is not a disease, potential anti-ageing drugs could be FDA approved separately for different conditions. The first study to test broad-spectrum protection capacity of metformin, the TAME study, is underway (Barzilai et al. 2016). In addition, it was shown that rapamycin/everolimus pre-treatment dramatically improves flu vaccination and immune response in the elderly (Mannick et al. 2018). In mice, it also lowers the incidence of tumours (Anisimov et al. 2011), and it shows promising results in the field of neurodegeneration (Bove et al. 2011). This supports the idea that targeting healthy ageing might have multiple beneficial outputs.

Our novel drug discovery platform will inform on new anti-ageing mechanisms, currently dominated by IIS and mTOR signalling pathways as well as dietary restriction regimes. Many drugs targeting growth pathways are already available from cancer research where they are used in very high doses. With the CellAgeClock, we examined which of these compounds could be disease preventative at very low concentrations. Our experimental setup is also suitable for nutraceutical approaches whereby dietary supplements can be rigorously tested for their effect on ageing. Overall, we expect our novel discovery platform to accelerate the discovery of strongly sought after anti-ageing drugs and geroprotective strategies to improve healthy human ageing.

## Materials and methods

### Cell culture and reagents

Normal finite lifespan human mammary fibroblasts (HMFs) were obtained from reduction mammoplasty tissue of a 16-year-old individual, donor 48 by Dr Martha Stampfer (University Berkeley) who has all required IRB approvals to distribute these cell samples and MTA agreement set in place with Dr Cleo Bishop laboratory. Independent cultures from these cells were serially passaged from passage 9 through to passage 20 and aliquots taken upon each passage for Illumina Infinium Methylation EPIC analysis. HMFs were maintained in Dulbecco’s Modified Eagles Medium (DMEM) (Life Technologies, UK) supplemented with 10% foetal bovine serum (FBS) (Labtech.com, UK), 2mM L-glutamine (Life Technologies, UK) and 10 μg/mL insulin from bovine pancreas (Sigma).

Normal finite lifespan human dermal fibroblasts (HDFs) were obtained from face lift dermis following a kind donation from an anonymous healthy patient under standard ethical practice, reference LREC No. 09/HO704/69. HDFs were grown in DMEM with 4 mM L-glutamine (Life Technologies), supplemented with 10% FBS.

Cells were plated at 7,500 cells/cm^2^ in T25 cell culture flask in 5ml of media to which 5μl of appropriate drug or vehicle control was added. Media was changed every two days and cells were passaged every 7 days and trypsinisation was used to detach the cells. All cells were routinely tested for mycoplasma and shown to be negative.

### Immunofluorescence microscopy and high content analysis

Cells were washed in phosphate buffered saline (PBS), fixed for 15 minutes with 3.7% paraformaldehyde with 5% sucrose, washed and permeabilised for 15 minutes using 0.1% Triton X in PBS (30 minutes for anti-nucleolin antibody) then washed and blocked in 0.25% bovine serum albumin (BSA) in PBS before primary antibody incubations. Primary antibodies used were anti-IL6 (R&D Systems, 1:100; overnight 4°C), anti-nucleolin (Santa Cruz, 1:2000, overnight room temperature), anti-p16 (Proteintech, 1:500, overnight 4°C), anti-p21 (12D1, Cell Signalling, 1:2000, overnight 4°C). Cells were incubated for 2 hours at room temperature with the appropriate AlexaFluor-488, AlexaFluor-546 or AlexaFluor-647 conjugated antibody (1:500, Invitrogen), DAPI (1:1,000 from 1mg/mL stock) and CellMask Orange or Deep Red (1:200,000, Invitrogen). Images were acquired using the IN Cell 2200 or 6000 automated microscope (GE) and HCA was performed using the IN Cell Developer software (GE).

### Z score generation

For each of the parameters analysed, significance was defined as one Z score from the negative control mean. Z scores were generated according to the formula below: Z score = (mean value of experimental condition − mean value of vehicle control/standard deviation (SD) for vehicle control.

### Senescence-associated beta-galactosidase (SA-β-Gal) assay

Cells were washed in PBS, fixed for 5 minutes with 0.2% glutaraldehyde, washed and incubated for 24 hour at 37°C (no CO_2_) with fresh senescence-associated beta-galactosidase (SA-β-Gal) solution: 1mg of 5-bromo-4-chloro-3-indoyl β-D-galactosidase (X-Gal) per mL (stock = 20mg of dimethylsulfoxide per ml) / 40mM citric acid/sodium phosphate, pH 6.0 / 5mM potassium ferrocyanide / 5mM potassium ferricyanide / 150mM NaCl / 2mM MgCl_2_). Cells were stained with Hoechst 33342 (1:10,000 from 10mg/mL stock) for 30 minutes. Images were acquired using the IN Cell 2200 automated microscope and HCA was performed using the IN Cell Developer software.

### Genomic DNA extraction

For isolation of Genomic DNA from primary human fibroblasts we used QIAamp DNA micro kit (56304) and we followed manufacturers protocol, with an additional washing steps with 500μl AW2 buffer and 500μl 80% ethanol to improve purity. DNA quantification and purity was determined by Nanodrop and QuBit. For bisulfite conversion EZ DNA methylation kit was used (D5001).

### Preparation of methylation array data

For each sample, 500ng high-quality DNA was bisulphite converted using the EZ DNA methylation kit (Zymo Research), using the alternative incubation conditions recommended for use with Illumina methylation arrays. Bisulphite converted DNA was eluted in 12ul elution buffer. Methylation was analysed using the Infinium Human Methylation EPIC array (Illumina) using standard operating procedures at the UCL Genomics facility. The EPIC array data have been deposited into ArrayExpress at the European Bioinformatics Institute (https://www.ebi.ac.uk/arrayexpress/) under accession number E-MTAB-8327.

### Pre-processing of methylation array data

DNA methylation array data was processed using the minfi package (Fortin et al. 2017) within R (R Core Team, 2013). Initial QC metrics from this package were used to remove low-quality samples. Probes were filtered using a detection p-value cut-off >0.01 and normalised using the Noob procedure. Cross-hybridising probes were removed from analysis based on the list published in McCartney et al. (McCartney et al. 2016). The training and test sets were pre-processed separately to obtain a fair estimate of the performance of the CellAgeClock.

### Estimation of sample age using existing epigenetic clocks

Following pre-processing of data, the epigenetic age of all samples was predicted using three epigenetic clocks; the Multi-tissue clock (Horvath 2013), the Skin and Blood clock (Horvath et al. 2018) and the PhenoAge clock (Levine et al. 2018) using the online DNA methylation calculator at http://dnamage.genetics.ucla.edu (Horvath 2013).

### Development of the CellAgeClock

The clock was built using a total of 39 samples, with six samples at each of the following passages; 10, 12, and 14; and seven samples at each of the following passages; 16, 18, and 20. This included both HDFs (n=12) and HMFs (n=27). 730,453 probes passed quality control measurements as described in the pre-processing section. A differential DNA methylation test was performed on this set of probes to identify CpGs undergoing significant DNA methylation changes with increasing cell passage. We used the linear regression approach for continuous variables implemented in minfi’s DMPFinder function (Fortin et al. 2017). This resulted in 2,543 differentially methylated CpGs at a p-value threshold of 1×10^−11^ which was selected using leave one out validation. Next, we built the clock model by elastic net regression using the DNA methylation levels of the 2,543 CpGs across all passages as input. We applied the glmnet function of the corresponding R package (Friedman et al. 2010) setting alpha to 0.5 and determining the lambda parameter by the internal cross validation function provided by glmnet. The elastic net regression model selected 42 CpGs as predictors of cell passage (Supplemental Table 1). We obtained the genomic annotation of these CpGs from Illumina’s EPIC manifest, and retrieved gene functions using DAVID (Huang da et al. 2009) where we selected the following sources for annotation: GOTERM_BP_DIRECT, GOTERM_CC_DIRECT, GOTERM_MF_DIRECT, ENTREZ_GENE_SUMMARY, OFFICIAL_GENE_SYMBOL and KEGG_PATHWAY.

We then tested the CellAgeClock on a different set of 26 samples, 22 HMFs and 4 HDFs, across the following passages: 9 (1 HMFs), 10 (2 HMFs), 11 (2 HMFs), 12 (1 HMFs), 13 (2 HMFs), 14 (2 HMFs), 15 (2 HMFs), 16 (4 HMFs and 2 HDFs), 18 (3 HMFs and 1 HDFs) and 20 (3 HMFs and 1 HDFs).

### Availability of the CellAgeClock

The CellAgeClock is available as a Jupyter Notebook in Python and can be retrieved from https://github.com/ucl-medical-genomics/CellAge-epigenetic-clock.

### Lifespan measurements

We used *white Dahomey* (*w*^*Dah*^) wild-type flies that were maintained and all experiments were conducted at 25°C. Flies were kept on a 12 h light:12 h dark cycle at constant humidity using standard sugar/yeast/agar (SYA) medium. For all experiments, flies were reared at standard larval density by transferring 18 *μ*l of egg suspension into SYA bottles. Eclosing adults were collected over a 12-h period and allowed to mate for 48 h before sorting into single sexes and placed in vials containing either control or experimental drug food. For lifespan assays, flies were reared at standard density and maintained at 15 flies per vial and we used holidic media recipe food for all longevity assays (Piper et al. 2014). Flies were transferred to fresh food vials every 2-3 days and scored for deaths. At least 150 flies were used for each lifespan experiment.

### Western blot measurements

Whole flies or human primary cell pellet was homogenised in 2x Laemmli loading sample buffer (100 mM Tris pH 6.8, 20% glycerol, 4% SDS; Bio-Rad) containing 50 mM DTT, protease inhibitor (cOmplete Mini EDTA-free; Roche) and phosphatase inhibitor (PhosSTOP EASYpack; Roche) cocktails. Extracts were cleared by centrifugation and approximately 20 μg of protein extract was loaded per lane on a polyacrylamide gel. Proteins were separated and transferred to nitrocellulose membrane. The following antibodies were used at the indicated dilutions: H3 (Cell Signaling Technology; 1:2000; 4499S), pS6K (Cell Signaling Technology; 1:1000; 9206S), total S6K (Santa Cruz; 1:1000; 8418), p4EBP (Cell Signalling Technology, 1:500; 2855S), non-phospho4E-BP (Cell Signalling Technology; 1:500; 4923S), pAkt (Cell Signalling; 1:1000; 4060), pAkt (Cell Signalling; 1:1000; 4056), total Akt (Cell Signalling; 1:1000; 9272), pERK (Cell Signalling; 1:1000; 4370), total (Cell Signalling; 1:1000; 4692). Blots were developed using the ECL detection system (GE, Amersham), and analysed using FIJI software (US National Institutes of Health). We used precasted TGX stain-free gels from Bio-Rad (567-8123 or 567-8124) according to the manufacturer’s instructions.

### Statistical analysis

Statistical analysis was performed using R. Log-rank tests were performed on lifespan curves.

## Online content

The CellAgeClock is available from GitHub at https://github.com/ucl-medical-genomics/CellAge-epigenetic-clock. All methylation microarray data reported in this study have been deposited in the ArrayExpress (https://www.ebi.ac.uk/arrayexpress/) public repository and are accessible under accession number E-MTAB-8327.

## Acknowledgements

IB acknowledges funding from ERC StG 311331, ERC PoC 842174, Royal Society Research Grant, The Bill Lyons foundation. SB acknowledges support from the Wellcome Trust (218274/Z/19/Z). This work was supported in part by the CRUK-UCL Centre Award [C416/A25145] awarded to IB and SB. EJT has been funded by MRC (MR/K501372/1) and BBSRC (BB/P002579/1) and DM is funded by the BBSRC (BB/N503629/1). JG, and MS are supported by U.S. Department of Energy under Contract No. DE-AC02-05CH11231 and the Congressionally Directed Medical Research Programs Breast Cancer Research Program Era of Hope Scholar Award BC141351.

## Authors contribution

SB, IB developed initial concept. SB, IB, RL and CLB finalised concept developing and designed experiments. RL, SE and APW analysed data. RL developed the CellAgeClock. CLB and EJT provided cell culture expertise. CL, EJT, ERS, VEMM, DM and IB performed all experiments. SE, SB, IB, RL, CLB wrote the manuscript. JCG and MRS provided critical reagents. All authors discussed results and commented on and approved the manuscript.

## Additional information

Correspondence and requests for materials should be addressed to IB, CLB, RL and SB.

## Supplemental Figure Legends

**Supplemental Figure 1.**
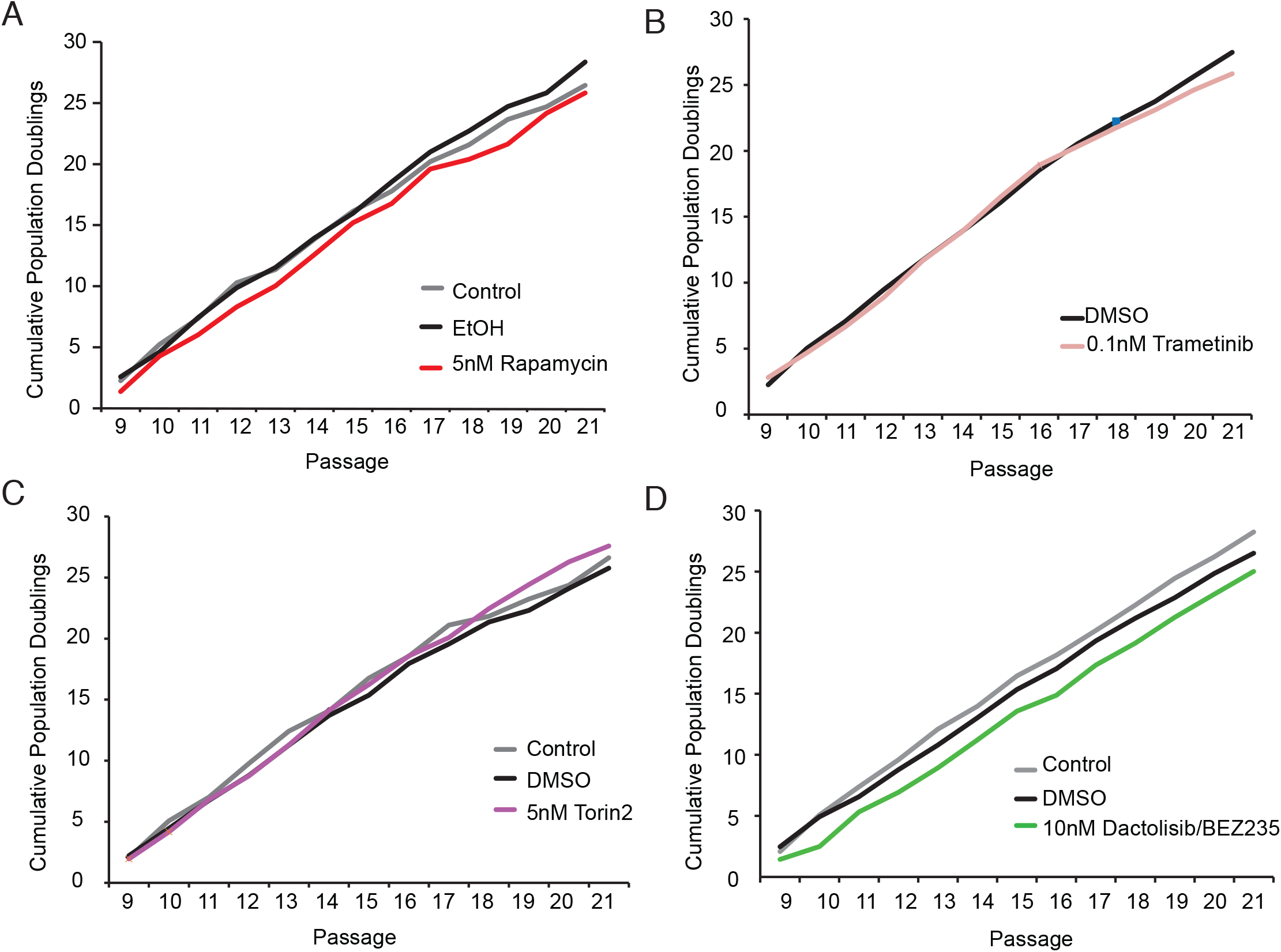
Drug treatment of HMF did not affect cell growth as measured by population doubling. Represented are population doubling measurements from passage 9 to passage 21 for A) rapamycin (5nM), B) trametinib (0.1nM), C) Dactolisib/BEZ235 (10nM), D) Torin2 (5nM) and corresponding vehicle controls.

**Supplemental Figure 2.**
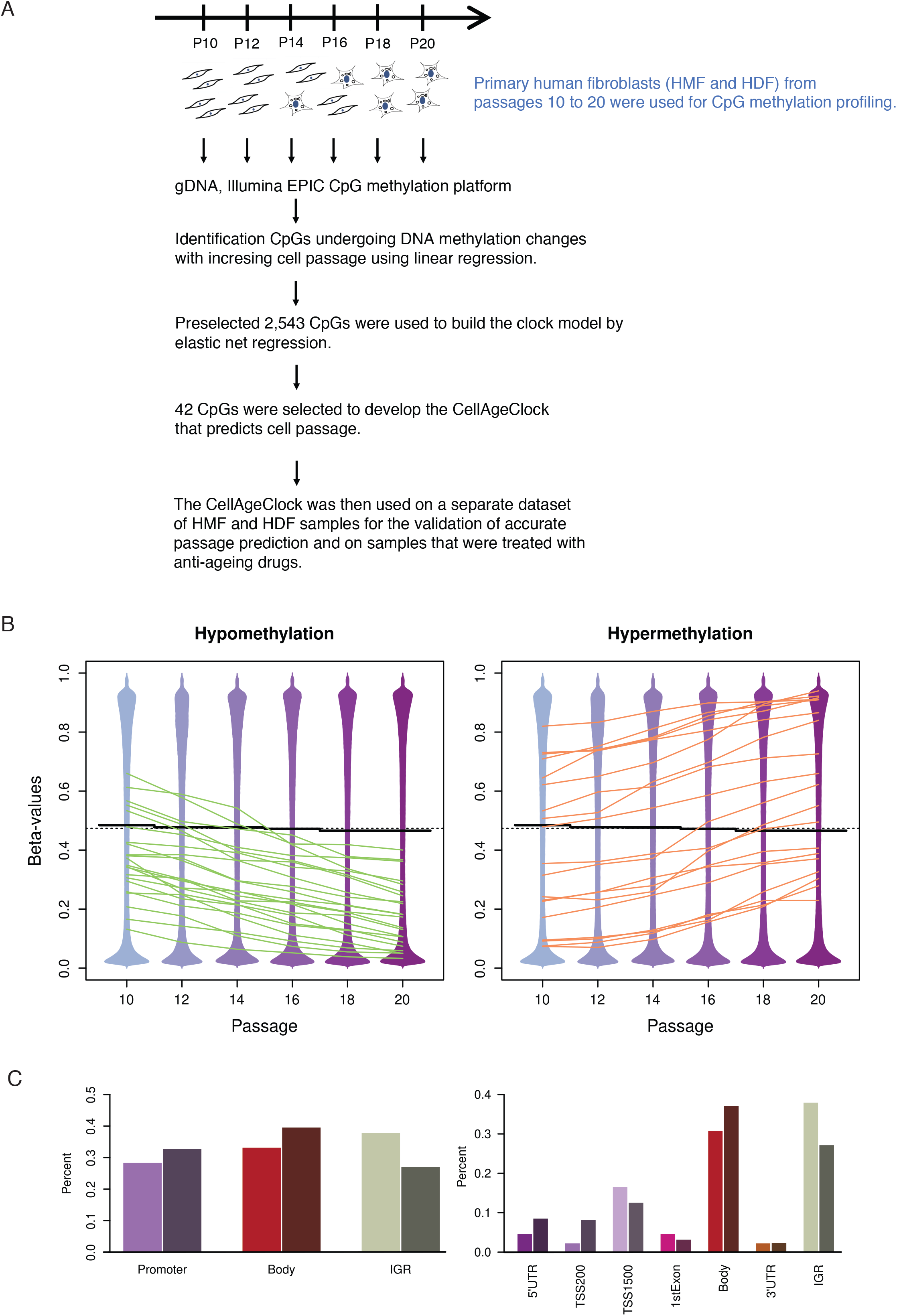
A) Scheme for developing of the CellAge clock. A) Primary human mammary fibroblasts and dermal fibroblasts, from passages 10 to 20, were used for CpG methylation profiling and using elastic net regression 42 CpGs were selected to develop the CellAgeClock that predicts cell passage. B) The beanplots showing the overall distribution of DNA methylation values (all CpGs, n=736,001) are presented. Overall DNA methylation levels decrease with increasing cell passage. The lines show where the clock CpGs sit in the distributions (mean per passage group), and are connected across passages to see how their methylation levels change. The plots show the mean methylation changes the clock CpGs are undergoing with increasing passage, separated by those that become hypo- and hypermethylated, in the contextof the global DNA methylation patterns of all CpGs. C) Barplots showing the percentage of CpGs falling into the corresponding genomic feature. Lighter bars show the percentages of clock CpGs (x% of the 42 CpGs). Darker bars show the background distribution (all other CpGs analyzed = 735,959 CpGs). Clock CpGs fall more often into IGRs. Barplots on the left are separated into gene promoters, gene bodies and IGRs for which in the CellAgeClock CpGs we found 12, 14 and 16, respectively. More detailed separation of the CellAgeClock CpGs into 5’UTR (2 CpGs), TSS200 (1 CpG), TSS1500 (7 CpGs), 1^st^ exon (2 CpGs), gene body (13 CpGs) and 3’UTR (1CpG) and IGR (16 CpG).

**Supplemental Figure 3.**
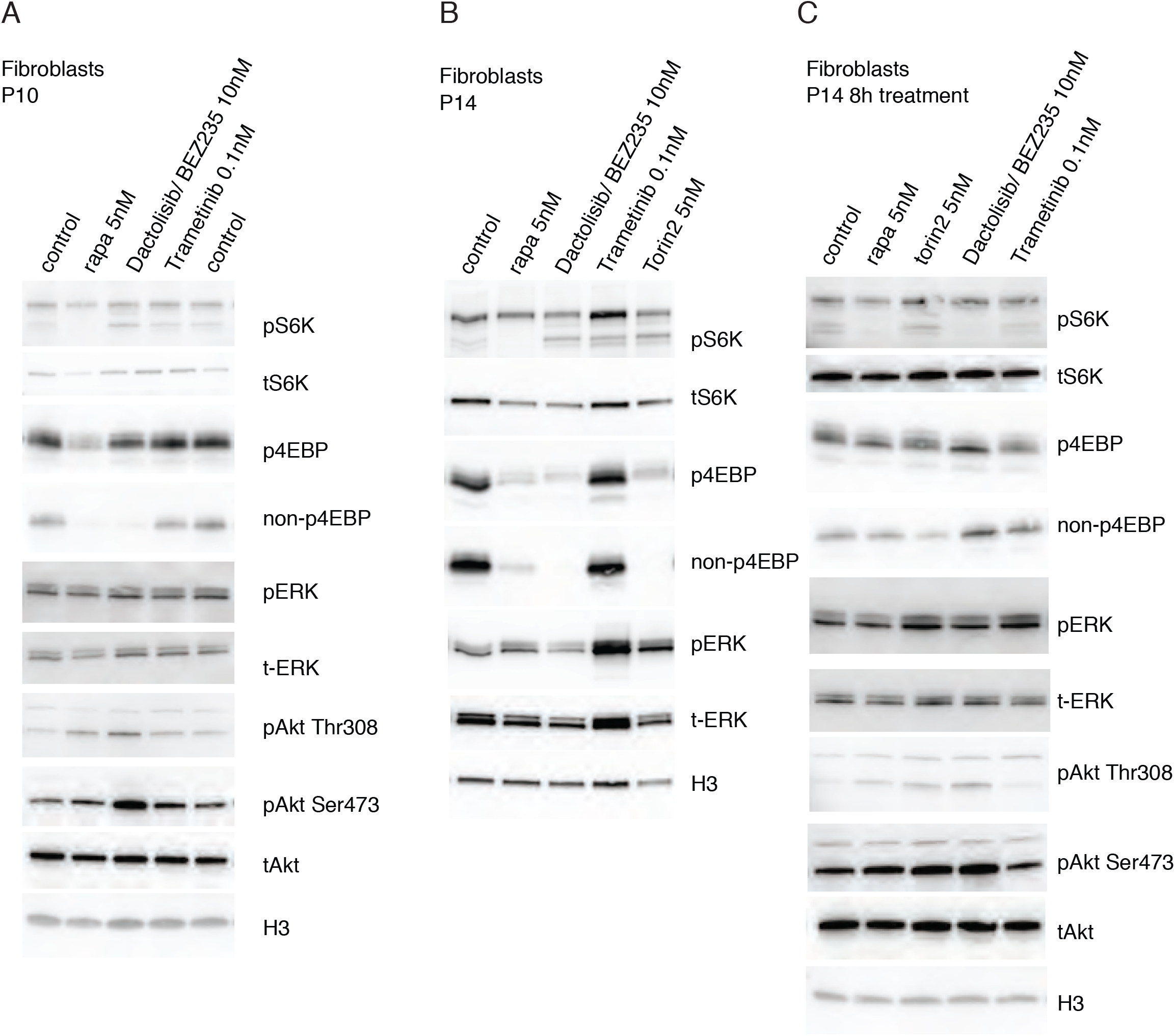
Western blot analysis on HMFs treated with anti-ageing drugs. Western blots were probed for phospho-S6K (Thr 389) showing p85 S6K Thr412 and lower band p70 S6K Thr389, total S6K, phospho-4EBP (Thr37/46), non-phospho-4EBP, pERK (Thr202/Tyr204), total ERK, and H3 as a loading control. Western blots analysis of A) HMF passage 10 and B) HMF passage 14 C) HMF passage 14 only 8h treatment with anti-ageing drugs. The analysis reflect rapid (S6K) and slower (4EBP) decreases in phosphorylation, increased phosphorylation (Akt Ser 473 and Akt Thr308) owing to S6K-IRS negative feedback loop, and resilience to phosphorylation changes owing to pMEK rebound and pERK feedback reactivation.

**Supplemental Figure 4.**
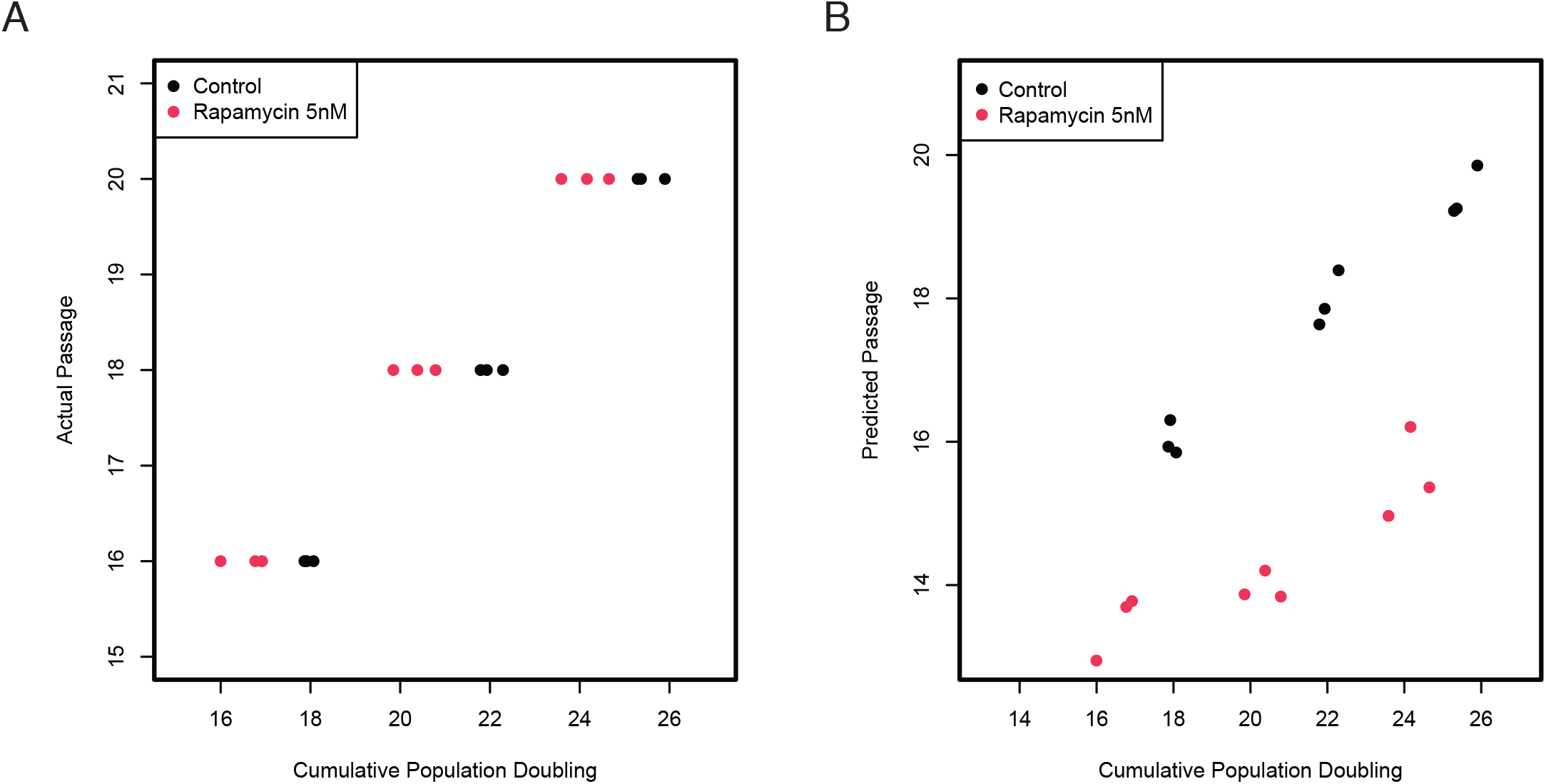
A) A scatterplot of the cumulative population doubling against the actual passage for control and rapamycin samples. B) A scatterplot of the cumulative population doubling against the predicted passage from CellAge for control and rapamycin samples.

**Supplemental Figure 5.**
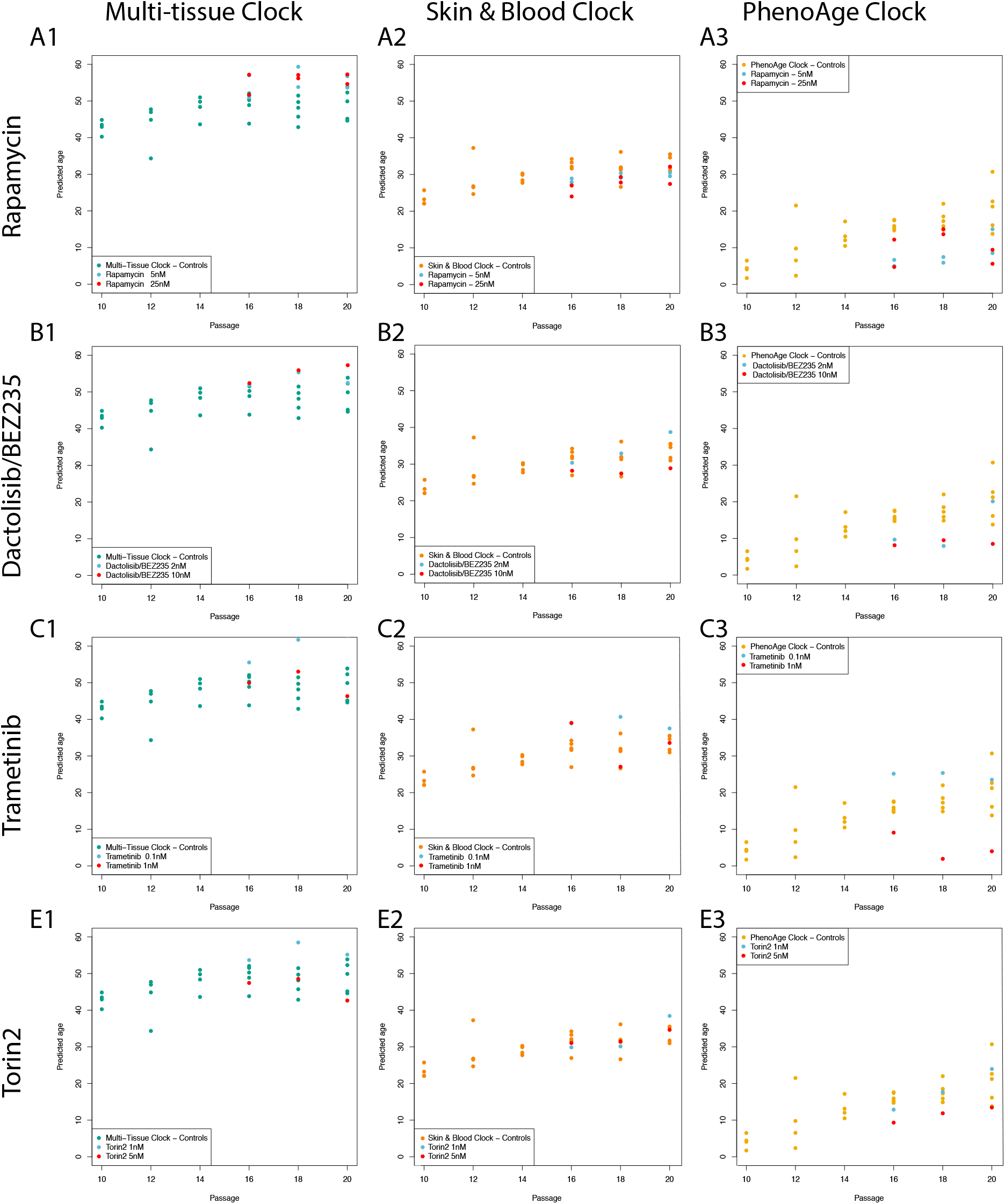
Predicted age of control samples and samples treated with potential anti-ageing drugs using three existing epigenetic clocks. Predictions of epigenetic age made using the Multi-Tissue (A1-D1, green), Skin & Blood (A2-D2, orange) and PhenoAge (A3-D3, yellow) clocks. Age predictions are indicated for samples treated with rapamycin (A1-A3), Dactolisib/BEZ235 (B1-B3), trametinib (C1-C3), torin2 (D1-D3), as compared with the set of control samples. These comparisons show that the existing clocks do not consistently indicate the anti-ageing effects of these treatments on cultured cells.

