## Supplemental Table 2 for "A CellAgeClock for expedited discovery of anti-ageing compounds"

***Supplemental Table 2:*** *Spearman's rank correlation and RMSE of the different clocks' predictions with actual cell passage of 26 samples that were not used to build the clock.*

| Clock | Spearman’s Rho | p-value | RMSE |
|---|---|---|---|
| Multi-tissue clock | 0.41 | 0.03837 | 33.21 |
| PhenoAge clock | 0.62 | 0.00070 | 14.01 |
| Skin and Blood clock | 0.81 | 4.8e-07 | 11.89 |
| CellAgeClock | 0.98 | < 2.2e-16 | 0.79 |
